# Analysis of 56K genomes identifies the relationship between antibiotic and metal resistance co-Occurrence and the spread of multidrug-resistant non-typhoidal Salmonella

**DOI:** 10.1101/2021.06.01.446601

**Authors:** Gavin J Fenske, Joy Scaria

**Affiliations:** Department of Veterinary and Biomedical Sciences, South Dakota State University, Brookings, South Dakota, USA, 57007

**Keywords:** Antibiotic Resistance, Metal Resistance, Co-Occurrence, Salmonella enterica, Genomics

## Abstract

*Salmonella enterica* is common foodborne pathogen that generates both enteric and systemic infections in hosts. Antibiotic resistance is common is certain serovars of the pathogen and of great concern to public health. Recent reports have documented the co-occurrence of metal resistance with antibiotic resistance in one serovar of *S. enterica*. Therefore, we sought to identify possible co-occurrence in a large genomic dataset. Genome assemblies of 56,348 strains of *S. enterica* comprising 20 major serovars were downloaded from NCBI. The downloaded assemblies were quality controlled and *in silico* serotyped to ensure consistency and avoid improper annotation from public databases. Metal and antibiotic resistance genes were identified in the genomes as well as plasmid replicons. Co-current genes were identified by constructing a co-occurrence matrix and group the genes using k-means clustering. Three groups of co-occurrent genes were identified using k-means clustering. Group 1 was comprised of the pco and sil operons that confer resistance to copper and silver respectively. Group 1 was distributed across four serovars. Group 2 contained the majority of the genes and little to no co-occurrence was observed. Metal and antibiotic co-occurrence was identified in group 3 that contained genes conferring resistance to: arsenic, mercury, beta-lactams, sulfonamides, and tetracyclines. Group 3 genes were also associated with an IncQ1 class plasmid replicon. Metal and antibiotic cooccurrence was isolated to one clade of *S. enterica* I 4,[5],12:i:.

## INTRODUCTION

Antibiotic resistant *Salmonella enterica subspecies enterica* is classified as a serious threat to public health by the Centers for Disease Control and Prevention in the United States [1]. *S. enterica* can rapidly disseminate antimicrobial resistance (AMR) genes horizontally, given the pathogen infects a wide range of hosts. Such a broad host range presents a unique challenge to public health. It has been demonstrated that sub-clinical doses of veterinary antibiotics in feed can promote the acquisition of resistance genes to clinically relevant antibiotics [2]. In this way, *S. enterica* acts as an antibiotic resistance bridge between agricultural and clinical settings. To combat the spread of AMR in bacteria, metals such as copper and zinc have been proposed and adopted by some producers as an alternative to antibiotics [3, 4]. One supplement, copper sulphate, has been proposed as a growth promoter in swine feed since at least 1961 [5]. The interest in metals as alternatives to antibiotics as growth promotors is obvious: metals are not antibiotics, thus curbing the spread of antibiotic resistance, copper is relatively inexpensive, and the growth promoting benefits, namely feed conversion efficiency, are retained when using the metal. When the European Union decided to ban the use of antibiotics as growth promoters in animal feed in 2006, many pig producers looked to copper sulphate as an alternative [6]. However, it has been shown that pharmacological doses of copper sulphate in pig feed increases antibiotic resistance in *Escherichia coli* [7]. Conjugative plasmids harboring resistance to macrolides, glycopeptides, and copper sulphate have been isolated from strains of *Enterococcus faecium* [8]. When metal and antibiotic resistance genes are constituents in the same linkage groups (plasmids, transposons, phages, etc) metals will co-select for antibiotic resistance genes [9–11]. It is becoming evident that metals are not a suitable alternative to antibiotics in agricultural settings and provide that same selective pressures as the antibiotics they are intended to replace [12].

Co-occurrence of metal and antibiotic resistance genes in S. enterica have been documented in *S. enterica* I 4,[5],12:i:- [13–16] and it has been suggested that copper resistance has allowed for expansion of the once rare serovar [16]. Given *S. enterica’s* unique ability to act as a zoonotic disseminator of AMR and metal resistance genes, we sought to identify co-occurrence between metal and antibiotic resistance genes in a query set of > 56,000 genomes comprising 20 major serovars. We specifically focused upon metal resistance genes that are plasmid bound to focus our analysis on genes that could be horizontally transmitted.

## RESULTS

### Broad screen for metal and antibiotic co-occurrence in S. enterica

The aim of our study was to identify co-occurrence between metal and antibiotic resistance genes in *S. enterica* with the potential for recombination (plasmid bound). To begin the screening, approximately 81,000 genome assemblies were downloaded from NCBI, quality checked, and in silico serotyped. The twenty most abundant serotypes, all containing greater than or nearly 1,000 assemblies (*S. enterica* Javiana, n = 995), were taken for further analysis (table 1).

**Table 1.** List of serovars and number of genome assemblies, median assembly length, and date range of assemblies used to identify co-occurrence of metal and antibiotic resistance.

| Serotype | Number of genomes | Median Length (Mb) | Date Range |
| --- | --- | --- | --- |
| Enteritidis | 11325 | 4.69 | 1950-2019 |
| Typhimurium | 8561 | 4.91 | 1958-2019 |
| Kentucky | 4131 | 4.92 | 1972-2019 |
| Infantis | 3866 | 4.94 | 1971-2019 |
| I 4,[5],12:i:- | 3657 | 4.95 | 1985-2019 |
| Newport | 3646 | 4.76 | 1975-2019 |
| Typhi | 2736 | 4.74 | 1958-2019 |
| Heidelberg | 2454 | 4.89 | 1979-2019 |
| Montevideo | 1841 | 4.65 | 1997-2019 |
| Agona | 1718 | 4.82 | 1952-2019 |
| Muenchen | 1641 | 4.79 | 1987-2019 |
| Saintpaul | 1570 | 4.79 | 1974-2019 |
| Anatum | 1560 | 4.73 | 1993-2019 |
| Senftenberg | 1329 | 4.81 | 2001-2019 |
| Schwarzengrund | 1130 | 4.81 | 2000-2019 |
| Mbandaka | 1083 | 4.75 | 2000-2019 |
| Braenderup | 1044 | 4.69 | 1999-2019 |
| Hadar | 1038 | 4.74 | 1988-2019 |
| Derby | 1023 | 4.87 | 1986-2019 |
| Javiana | 995 | 4.61 | 1995-2019 |

The final dataset totaled 56,348 *S. enterica* assemblies. Metal resistance, antibiotic resistance, and plasmid replicons were predicted from each assembly. To gain understanding into the temporal distribution of metal and antibiotic resistance, the mean number of antibiotic and metal resistance genes were plotted against year of collection as shown in Figure 1. Both resistance types show serovar specific trends. Many of the serovars show no trend between mean number of antibiotic and metal resistance genes over time. However, three serovars, *S. enterica* Kentucky, *S. enterica* I 4,[5],12:i:-, and *S. enterica* Infantis, have increasing antibiotic and metal resistance genes over time. The trend is most evident in *S. enterica* I 4,[5],12:i:-, with the mean number of metal resistance genes increasing nearly 3 fold in 15 years. S. enterica Schwarzengrund and S. enterica Senftenberg showed constant metal resistance with no trend observed to antibiotic resistance. S. enterica Enteritidis, the most abundant serovar, showed almost no antibiotic nor metal resistance homologues.

**Figure 1.**
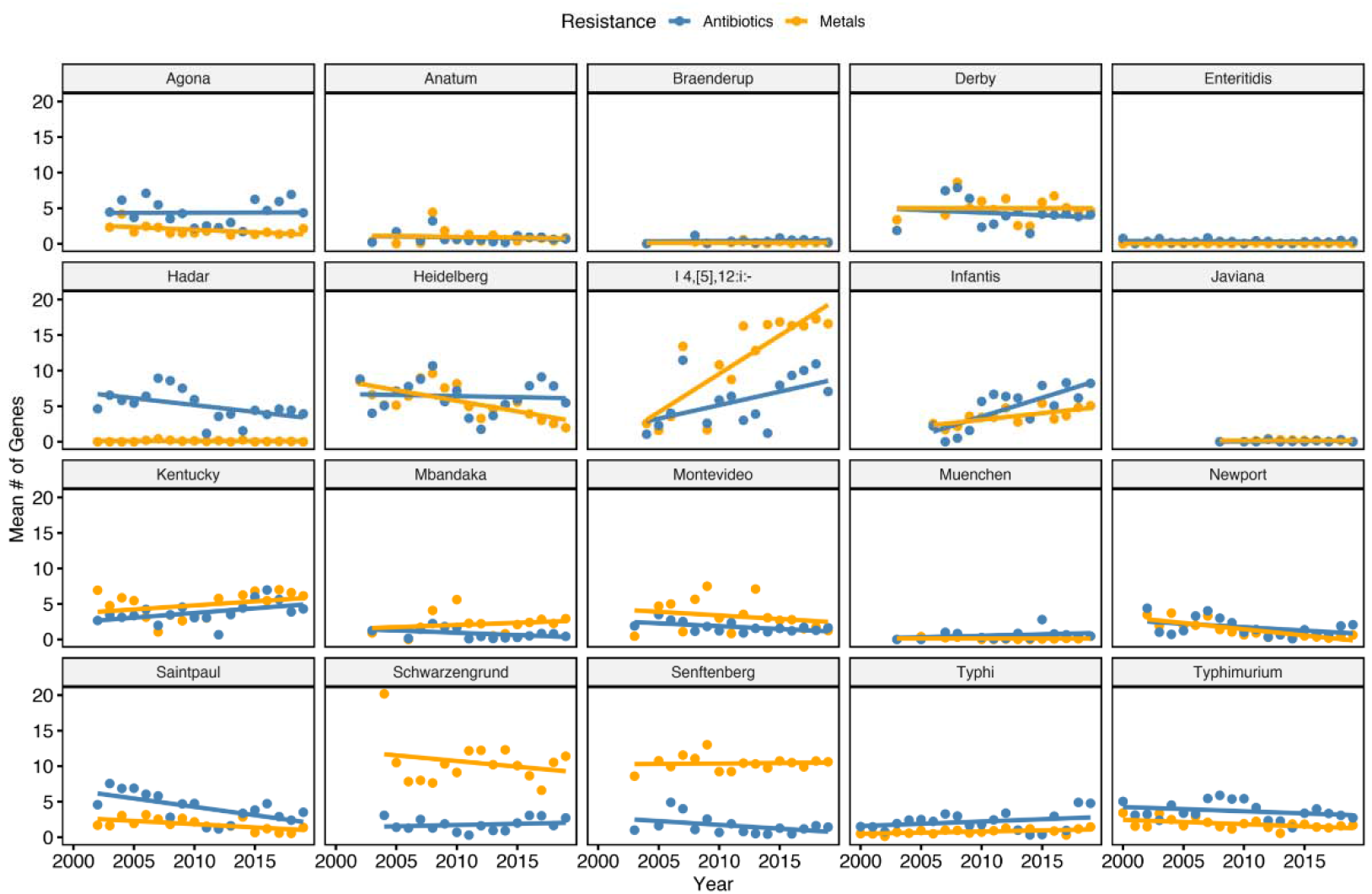
Temporal distribution of antibiotic and metal resistance genes by S. enterica serovar. The mean number of antibiotic and resistance homologues identified in each serovar is plotted against the date of collection. Only dates that contained n > 10 genomes were considered to control for outliers.

### Co-occurrence of metal and antibiotic resistance

Co-occurrence between plasmid replicons (included as the focus was to identify possible recombinant resistance), metal resistance genes, and antibiotic resistance genes was identified by applying k-means clustering to a co-occurrence matrix (see methods). Three groups of co-occurrent genes were identified (figure 2A). To better visualize the groups identified by k-means clustering, the co-occurrence network was plotted, and groups were colored respective to figure 2A (figure 2B). Group 1 was the smallest cluster set containing only 11 genes comprising the *pco* and *sil* operons. Said operons confer resistance to copper and silver respectively. Group 2 represents most plasmid replicons, metal resistance genes, and antibiotic resistance genes. Little co-occurrence is observed in group 2 save for two plasmid replicons (IncFIB(S)_1 and IncFII(S)_1). The small node sizes of other constituents in group 2 suggests that gene turnover may be high in the associated plasmids. The majority of antibiotic and metal resistance genes are not co-occurrent in the dataset of *S. enterica* genomes. However, co-occurrence was identified in group 3. Homologues in group 3 confer resistance to metals arsenic and mercury, antibiotic classes of beta-lactams, sulfonamides, tetracyclines, and contains one plasmid replicon: IncQ1_1. The top down approach and simple k-means clustering was able to identify that co-occurrence between metal and antibiotic resistance is occurring in some genomes of S. enterica. However, it also showed that the effect is only occurring in a small subset of genes, and that most are not carried together. The full list of homologues in groups 1 and 3 are presented in table 2.

**Figure 2.**
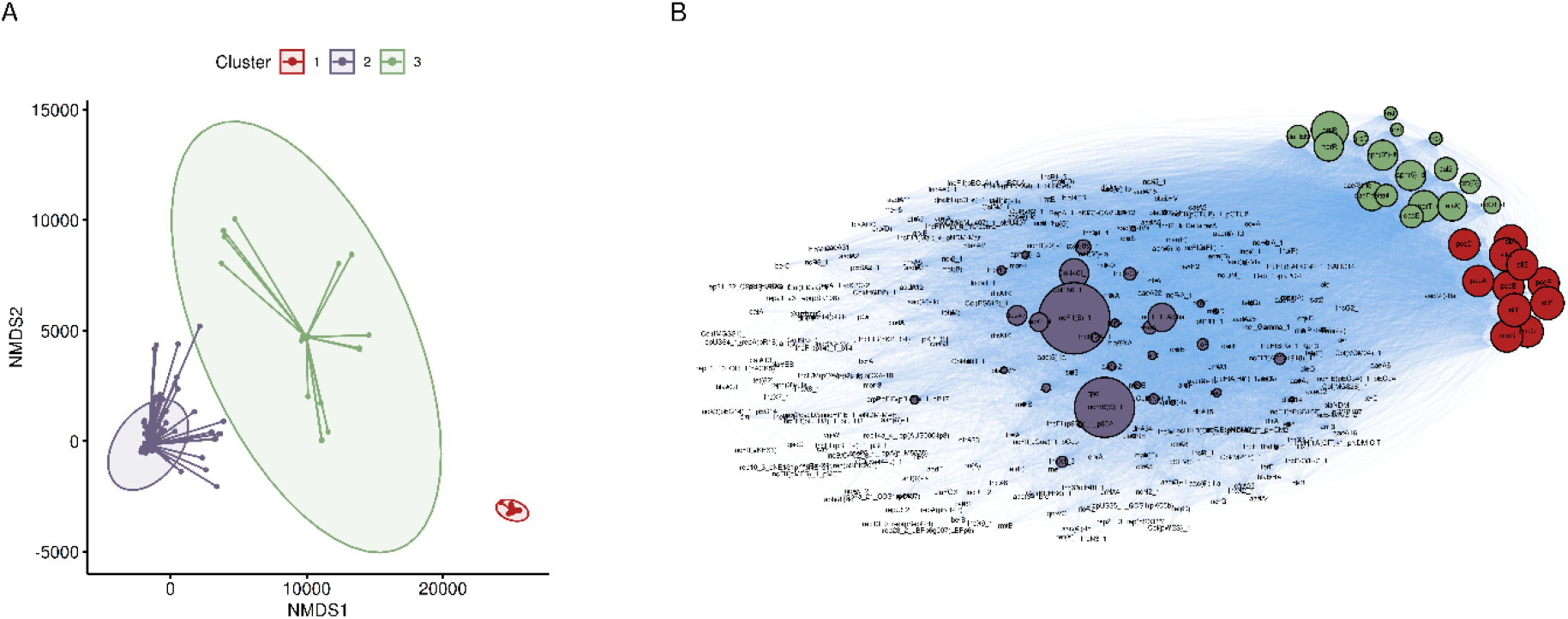
Co-occurrent gene clusters identified in S. enterica genomes. (A) Non-metric Multi-dimensional Scaling plot of a co-occurrence matrix generated from plasmid replicons, antibiotic and metal resistance homologues. K-means clustering was used to group genes. An elbow plot was used to pick the optimal number of clusters. Clusters are drawn as a star plot with lines connecting each gene to the center of the cluster. Gaussian distributions were drawn around each cluster. (B) Network representation of co-occurrence network. Nodes are scaled to edge weight (number of connections) and are colored consistent to A.

**Table 2.** Major groups of co-occurrent genes identified by k-means clustering.

| Cluster | Gene | Class | Subclass |
| --- | --- | --- | --- |
| 1 | pcoA | Copper | Copper |
|  | pcoB | Copper | Copper |
|  | pcoC | Copper | Copper |
|  | pcoD | Copper | Copper |
|  | pcoR | Copper | Copper |
|  | pcoS | Copper | Copper |
|  | silA | Silver | Silver |
|  | silB | Silver | Silver |
|  | silC | Silver | Silver |
|  | silE | Silver | Silver |
|  | silF | Silver | Silver |
| 3 | aph(3'')-Ib | Aminoglycoside | Streptomycin |
|  | aph(6)-Id | Aminoglycoside | Streptomycin |
|  | arsA | Arsenic | Arsenite |
|  | arsB | Arsenic | Arsenite |
|  | arsC | Arsenic | Arsenate |
|  | arsD | Arsenic | Arsenite |
|  | arsR | Arsenic | Arsenite |
|  | blaTEM | Beta-Lactam | Beta-Lactam |
|  | IncQ1_1 | Plasmid Replicon | NA |
|  | merR | Mercury | Mercury |
|  | merT | Mercury | Mercury |
|  | qacE | Quaternary Ammonium | Quaternary Ammonium |
|  | qacEdelta1 | Quaternary Ammonium | Quaternary Ammonium |
|  | sul1 | Sulfonamide | Sulfonamide |
|  | sul2 | Sulfonamide | Sulfonamide |
|  | tet(A) | Tetracycline | Tetracycline |
|  | tet(B) | Tetracycline | Tetracycline |

To better visualize the distribution of group 1 and 3 gene clusters (moderate to high co-occurrent genes), a presence-absence matrix of group 1 and 3 genes was plotted against the phylogeny of *S. enterica* I 4,[5],12:i:-, S. enterica Infantis, S. enterica Kentucky, S. enterica Schwarzengrund, and S. enterica Senftenberg (serovars that showed trends in figure 1) as shown in figure 3. Group 1 (sil and pco) genes are broadly distributed among S. enterica I 4,[5],12:i:-, S. enterica Kentucky, S. enterica Schwarzengrund, and S. enterica Senftenberg, but are largely absent from S. enterica Infantis. However, group 1 genes only confer resistance to the metals silver and copper, and not antibiotics. Only *S. enterica* I 4,[5],12:i:-assemblies consistently contained loci from group 3, the co-occurrent group of metal and antibiotic resistance genes. Therefore, we decided to focus the study upon *S. enterica* I 4,[5],12:i:- and the possible mechanisms of the co-occurrence. Recent research has proposed the addition of metals, namely silver and copper, to medical devices, food pack

**Figure 3.**
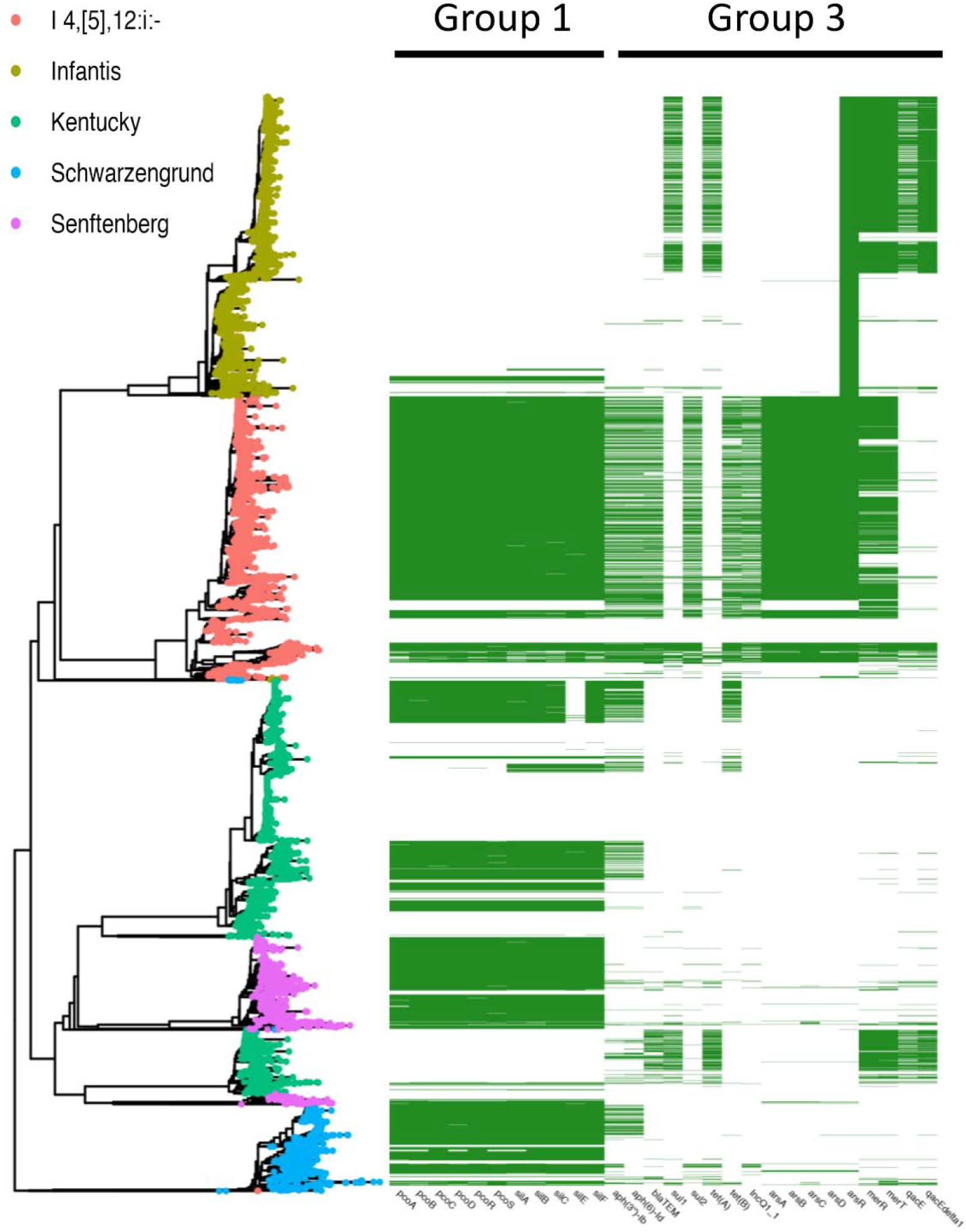
Distribution of group 1 and 3 genes among *S. enterica* I 4,[5],12:i:-, S. enterica Infantis, S. enterica Kentucky, S. enterica Schwarzengrund, and S. enterica Senftenberg. A prescence-absence matrix of group 1 and 3 genes is plotted against a neighbor-joining tree constructed from pairwise distance matrix. Group 1 (pco and sil operons) is found in S. enterica Kentucky, S. enterica Schwarzengrund, and S. enterica Senftenberg but is mostly absent from S. enterica Infantis. Group 3, which contains metal and antibiotic resistance genes, is almost exclusive to *S. enterica* I 4,[5],12:i:-.

### *S. enterica* I 4,[5],12:i:-

*S. enterica* I 4,[5],12:i:-was the only serovar to contain group 3 genes in a substantive manner. The initial dataset was compiled from assembled genomes in order to facilitate speed. However, the differing assembly mechanisms and lack of SNP based phylogeny limits the accuracy. To remedy this, 1,455 sequencing reads of US S. enterica I 4,[5],12:i:-, 1,447 sequencing reads of US S. enterica Typhimurium and 329 sequencing reads of European S. enterica I 4,[5],12:i:-were compiled into a SNP phylogeny against a reference S. typhimurium DT104 assembly. The addition of the new data set allows for a validation step against the initial screening. Plasmid replicons, metal and antibiotic resistance were annotated in the second dataset exactly following the protocol for the large query set. A presence-absence matrix describing group 1, 3, and major plasmid replicons is plotted against the phylogeny of S. enterica I 4,[5],12:i:- and S. enterica Typhimurium in figure 4.

**Figure 4.**
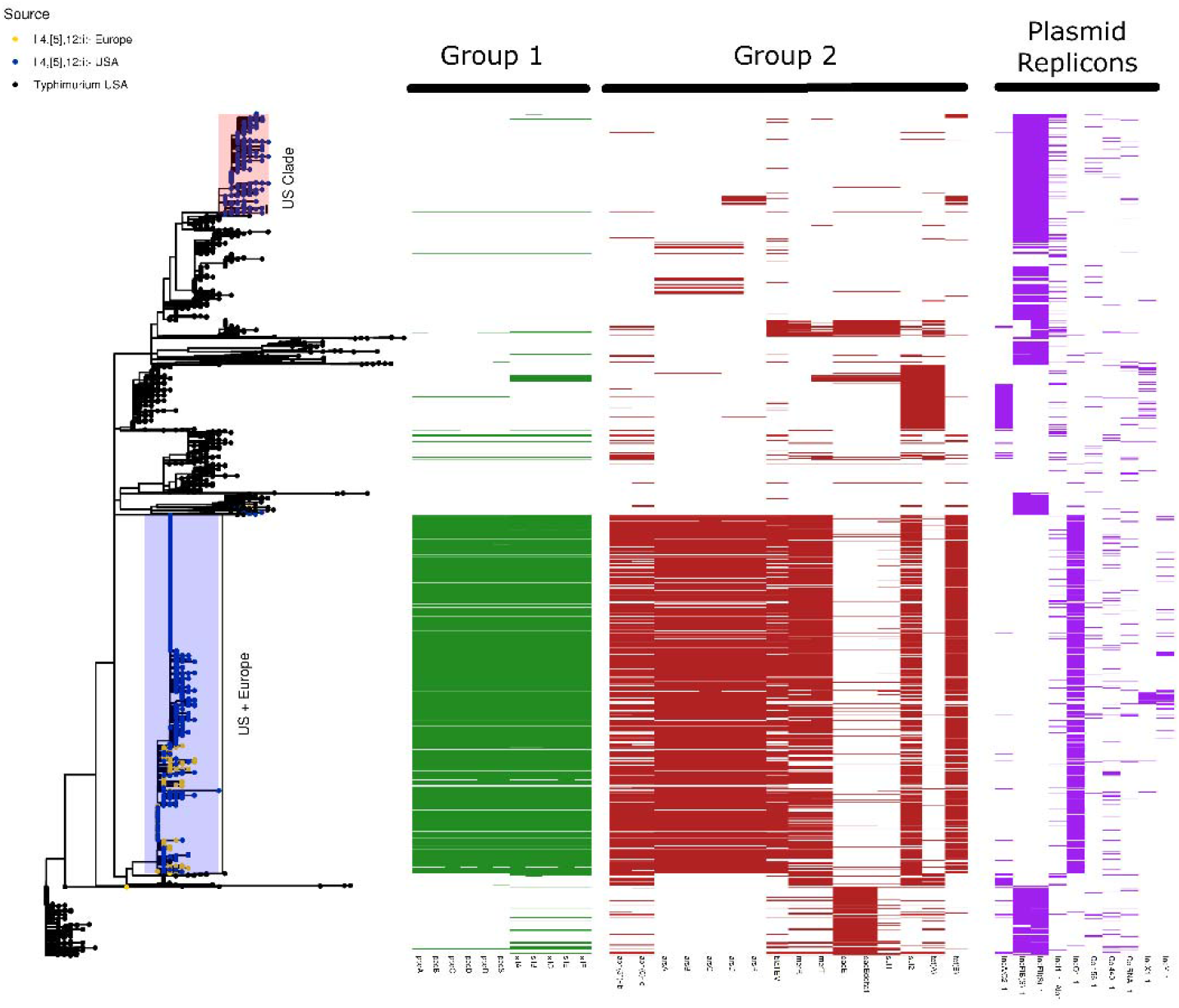
Distribution of group 1, group3, and major plasmid replicons in *S. enterica* I 4,[5],12:i:-. A presence-absence matrix for each gene group is plotted against a maximum-likelihood tree of S. enterica Typhimurium and *S. enterica* I 4,[5],12:i:-rooted a reference S. enterica Typhimurium DT104 assembly. Two major clades of *S. enterica* I 4,[5],12:i:- are observed, however only the clade containing both US and European isolates contains both group 1 and group 3 genes as well as the IncQ1_1 plasmid replicon.

Confirming the initial screening, group 1 and 3 genes were carried in S. enterica I 4,[5],12:i:-but were mostly absent from S. Typhimurium. Two major clades of S. enterica I 4,[5],12:i:- are observed: one comprising solely of US isolates and a mixed clade of US and European genomes. Notably, the mixed clade of US and European isolates contain both group 1 and 3 as well as the IncQ1_1 plasmid replicon. The S. enterica I 4,[5],12:i:-clade comprised solely of US genomes did not contain group 1 and 3 genes. To this end, the co-occurrence of metal and antibiotic resistance is not widespread throughout S. enterica, but rather is localized to one clade of S. enterica I 4,[5],12:i:-. The metadata for the detailed analysis of S. enterica I 4,[5],12:i:- and S. enterica Typhimurium is provided in supplemental table 5.

## DISCUSSION

In this work, we identified co-occurrence of metal and antibiotics in *S. enterica* I 4,[5],12:i:-by screening a broad set of 20 serovars totaling more than 56,000 genomes. The topdown approach allows for a non-biased and comprehensive approach compared to starting with metal resistant isolates and working upwards towards the serovar level. Ultimately, our hypothesis was incorrect as co-occurrence between metal and antibiotic resistance is not widespread in S. enterica. Rather, the phenomenon is limited to one clade of *S. enterica* I 4,[5],12:i:-. This result is interesting in that it highlights how the acquisition of resistance genes can lead to a rapid expansion of bacterial strains. *S. enterica* I 4,[5],12:i:- was rarely isolated in the 1990’s but was the third most isolated serovar of S. enterica in the European Union in 2017 [29] and the fifth most isolated serovar in the US in 2015 [30]. It has been suggested that the introduction of copper sulfate into pig feed, and the resistance to copper harbored by *S. enterica* I 4,[5],12:i:-strains, may have contributed to the rapid increase in prevalence [16]. *S. enterica* I 4,[5],12:i:-provides a meaningful example of the link of agricultural processes and human health. While many of the isolates did not contain the co-occurrence between antibiotics and metals we sought to identify, copper and silver resistance was identified in S. enterica Kentucky, enterica Schwarzengrund, and S. enterica Senftenberg.

The pco and sil operons are associated in S. enterica with Salmonella Genomic Island 4 (SGI-4). SGI-4 is a chromosomal island containing the *pco* and *sil* operons that may be excised from the chromosome by mitomycin C, and oxygen tension related stress [13]. SGI-4 was first designated in 2016 [16](see addendum) and is largely associated with *S. enterica* I 4,[5],12:i:- [13]. Indeed it has been stated that SGI-4 has only been discovered in I 4,[5],12:i:- [31]. Our results indicate the genomic island may be more widespread than believed. As metal use increases in medical and agricultural settings, the opportunity exists for metal resistant strains to rapidly expand in prevalence as has been witnessed with *S. enterica* I 4,[5],12:i:-. Indeed, a burn ward identified S. enterica Senftenberg as a causative agent of an outbreak in severe burn patients [32]. The isolated strains were resistant to the silver sulfadiazine that was applied daily to the burn patients. Another concern is the increasing use of copper sulphate as a growth promoter in chickens [33, 34]. S. enterica Schwarzengrund is commonly isolated from chickens and global outbreaks of multi-drug resistant strains have occurred [35]. If metal use in poultry agriculture continues, S. enterica Schwarzengrund outbreaks have the potential to increase in the coming years.

The serovar specific nature of group 3 genes (metal and antibiotic co-occurrence) is consistent with earlier reports [14, 36, 37]. Previous work has demonstrated that a clade *S. enterica* I 4,[5],12:i:-circulating the US is descendent from multi-drug resistant strains in Europe [38]. The inclusion of the IncQ1_1 plasmid replicon reinforces the claim as the gene was the most identified plasmid replicon of *S. enterica* I 4,[5],12:i:-in Italian isolates [14]. A plasmid harboring resistance to metals and antibiotics has been documented in an Australian *S. enterica* I 4,[5],12:i:-isolated from pig feces [15]. However, the plasmid was large (~ 275 kb) and annotated as a IncHI2 class plasmid, not an IncQ class as we report here. Additionally, it was found that the plasmid was able to conjugate to other species [15]. IncQ plasmids are unable to self-transfer and rely upon “helper plasmids” for transmission between bacterial cells [39]. The lack of other plasmid replicons in large numbers in the mixed US and European clade of *S. enterica* I 4,[5],12:i:-, and isolation of group 3 genes in the clade, suggests the plasmid and group 3 genes are being transmitted vertically rather than horizontally. Furthermore, IncQ class plasmids are evolutionary stable with one study documenting the presence of the plasmid in the environment over a 30 year period [40]. Taken together, we propose that the method of metal and antibiotic co-occurrence in the US and European monophasic strains studied here differs from the large IncHI2 plasmid isolated from an Australian strain of *S. enterica* I 4,[5],12:i:-. Complicating the narrative is the difficulty in accurately constructing plasmid contigs from shortread sequencing [41]. Isolating plasmids from a large number of *S. enterica* I 4,[5],12:i:-containing the IncQ1_1 and group 3 genes and sequencing with a long-read platform such as PacBio is needed to confirm our results.

## MATERIALS AND METHODS

### Salmonella enterica genome assembly acquisition

Genome assemblies and GenBank files for *S. enterica* isolates were downloaded from Genome on NCBI (ftp.ncbi.nih.gov::genomes/all/GCA/, October 10, 2019). Resultant GenBank files were parsed and only assemblies containing a collection date were included for further analysis. Downloaded assemblies were removed from the data set if the contig number was greater than 300. Genome assemblies were in silico serotyped using SISTR [17]. This was done to ensure consistent serovar annotation and limit the impact of improper or mislabeled serotypes. The top 20 serovars in the dataset were used for co-occurrence identification. 20 serovars were chosen as each contained greater than 1,000 genomes, save for S. enterica Javiana, which contained n = 995.

### Identification of plasmid replicons, metal and antibiotic resistance homologues

Plasmid replicons were identified in genome assemblies using the tool Abricate [18] against the PlasmidFinder [19] database. Hits were defined as sequences with >= 90% percent identity and >= 60% query coverage.

Open-reading frames (ORF) in assemblies were predicted using Prodigal [20]. Metal resistance homologues were identified in the resultant ORF files using GhostZ [21] to compare the ORF against the experimentally verified BacMet database (version 2.0) [22]. ORF were considered homologues when: e-value <= 1 E-6, percent identity >= 90 % and percent coverage >= 60%. As we aimed to identify metal resistance under possible recombination, only homologues marked “PLASMID” in the BacMet database were considered.

Antibiotic resistance homologues were identified in an analogous manner to the metal resistance genes, however, ORF were queried against the NCBI antimicrobial resistance database (BioProject: PRJNA313047) using the same parameters.

### Co-Occurrence identification

Plasmid replicons, metal and antibiotic resistance homologues from each assembly were compiled into a binary matrix. A co-occurrence matrix was generated by taking the cross product of the original binary matrix. Simply put, if two genes are contained in the same assembly, they yield a value of 1. If genes are contained across multiple assemblies together, the number will increase accordingly. To identify possible co-occurrence between genes, the co-occurrence matrix was subjected to k-means clustering using clusters. The optimal number of clusters was determined by constructing an “elbow plot” (the sum of square distances against k number of clusters). Matrix compilation and k-means clustering were conducted using R (R-core team). Igraph [23] was used to visualize the co-occurrence network. Nodes are scaled to edge weight number (larger nodes contain more co-occurrences).

### Phylogeny and S. enterica I,4,[5],12:i:- analysis

To compile a phylogeny estimation of 56,348 *S. enterica* genomes, MASH [24] sketches were drawn from each sample (10,000 sketches, kmer size of 21) and a pairwise distance matrix was generated. A neighbor-joining tree was constructed from the distance matrix using QuickTree [25].

Reads of European and US *S. enterica* I,4,[5],12:i:- and US *S. enterica* typhimurium and were downloaded from SRA hosted at NCBI (https://www.ncbi.nlm.nih.gov/sra). Resultant reads were mapped reads against a *S. enterica* Typhimurium DT104 NCTC 13348 reference strain downloaded from https://www.sanger.ac.uk/resources/downloads/bacteria/salmonella.html. Mapping and variant calling was conducted using Snippy [26] and only core variants were considered. A maximum likelihood from the SNP alignment was generated using IQTree [27] with a GTR+G4 substitution model and 1000 bootstraps. All phylogenies were visualized using the R package ggtree [28].

Reads were assembled into contigs using the package Shovill (https://github.com/tseemann/shovill) with a minimum contig length of 200 base pairs. Prodigal was used to predict ORF and plamid replicons, antibiotic and metal resistance genes were predicted in the same manner as the genomes of the large test set.

## Data Availability Statement

Salmonella genome data used in this analysis is public data already deposited in NCBI SRA. All salmonella genomes available in NCBI SRA up to October 10, 2019 was used for the analysis.

## Acknowledgments

Authors gratefully thank Dr. Marc W. Allard, and Dr. Ruth Timme, FDA CFSAN ORS and the GenomeTrakr Network labs which sequenced and submitted the Salmonella genome data to NCBI SRA which was used for the analysis in this manuscript.

